# Prediction and Curation of Missing Biomedical Identifier Mappings with Biomappings

**DOI:** 10.1101/2022.11.29.518386

**Authors:** Charles Tapley Hoyt, Amelia L. Hoyt, Benjamin M. Gyori

## Abstract

**Motivation:** Biomedical identifier resources (ontologies, taxonomies, controlled vocabularies) commonly overlap in scope and contain equivalent entries under different identifiers. Maintaining mappings for these relationships is crucial for interoperability and the integration of data and knowledge. However, there are substantial gaps in available mappings motivating their semi-automated curation.

**Results:** Biomappings implements a curation cycle workflow for missing mappings which combines automated prediction with human-in-the-loop curation. It supports multiple prediction approaches and provides a web-based user interface for reviewing predicted mappings for correctness, combined with automated consistency checking. Predicted and curated mappings are made available in public, version-controlled resource files on GitHub. Biomappings currently makes available 8,560 curated mappings and 41,178 predicted ones, providing previously missing mappings between widely used resources covering small molecules, cell lines, diseases and other concepts. We demonstrate the value of Biomappings on case studies involving predicting and curating missing mappings among cancer cell lines as well as small molecules tested in clinical trials. We also present how previously missing mappings curated using Biomappings were contributed back to multiple widely used community ontologies.

**Availability:** The data and code are available under the CC0 and MIT licenses at https://github.com/biopragmatics/biomappings.

## 1 Introduction

Standardizing the identification of small molecules, proteins, and other biomedical entities is an important step in creating and maintaining Findable, Accessible, Interoperable, and Reusable (FAIR) (Wilkinson *et al*., 2016) data in the life sciences. Resources that catalog and provide identifiers for such entities are essential for this effort. Such *identifier resources* include ontologies, taxonomies, and other controlled vocabularies, e.g., the Human Disease Ontology (DO) (Schriml *et al*., 2021), Medical Subject Headings (MeSH) (Rogers, 1963), and the Chemical Entities of Biomedical Interest (ChEBI) (Hastings *et al*., 2016). However, many identifier resources overlap in scope and include equivalent entities with different identifiers. For example, the small molecule cyclin-dependent kinase inhibitor *alsterpaullone* appears in several chemical- and drug-related identifier resources including ChEBI as `chebi:138488` and MeSH as `mesh:C120793`. Merging equivalent entities is crucial for tasks dependent on data integration like ontology merging (Geleta *et al*., 2022; Guo *et al*., 2022; Lambrix and Tan, 2008), entity linking (Gyori *et al*., 2022), construction of knowledge graphs (Himmelstein *et al*., 2017; Friedrichs, 2021; Nicholson and Greene, 2020), and automated systems biology model assembly (Gyori *et al*., 2017; Bachman *et al*., 2022). More generally, mapping between equivalent identifiers from different identifier resources is a ubiquitous task across computational life science analyses, workflows, and tools.

A *mapping* (often referred to as *ontology mapping* or *semantic mapping*) represents a relationship between two entities in different identifier resources using a specific predicate such as one representing an exact match (i.e., when the entities can be used interchangeably), a broad match (i.e., when one entity is a super-class of the other), or a narrow match (i.e., when one entity is a subclass of the other). Mappings can also carry additional metadata representing provenance for its creation. We refer to Figure 2 of Matentzoglu *et al*. (2022a) for a more detailed description of mappings. High-quality, semantically rich equivalence mappings are required to support merging and converting between identifiers from different resources. Therefore, many identifier resources provide equivalence mappings to one or more other resources. We refer to mappings provided directly by an identifier resource as *primary mappings*. Such mappings are typically curated by the maintainers of the resource and are provided along with entries in the resource in the form of cross-references. For example, the HUGO Gene Nomenclature Consortium (HGNC) (Yates *et al*., 2017) provides mappings from its identifiers for human genes to the Entrez gene database (Maglott *et al*., 2011), Ensembl (Zerbino *et al*., 2018), and several other identifier resources. Similarly, ontologies like the Monarch Disease Ontology (MONDO) (Vasilevsky *et al*., 2022) curate and distribute mappings to other identifier resources (e.g., DO, MeSH).

Though primary mappings from identifier resources are often available, a survey from Laadhar *et al*. (2020) of mappings in life science ontologies highlights several widespread issues such as the use of unspecific predicates (e.g., `oboInOwl:hasDbXRef`), the lack of standardization of the syntax and semantics of the targets of mappings, and a lack of detailed provenance metadata. The Simple Standard for Sharing Ontological Mappings (SSSOM) (Matentzoglu *et al*., 2022a) was recently developed to provide a standardized format for mappings and thereby increase their availability and interoperability. While SSSOM supports disseminating mappings through a common standard, it does not in itself provide a solution to identifying and curating missing mappings.

Several services aggregate, process, and redistribute mappings including BridgeDB (van Iersel *et al*., 2010), TogoID (Ikeda *et al*., 2022), and the Ontology Mapping Service (EMBL-EBI, 2022). However, these services only draw from primary mappings provided by identifier resources and are therefore unable to address gaps, lack of specificity, and lack of rich provenance metadata in these resources.

Despite existing curation and aggregation efforts, there still exist gaps in mappings between major resources (e.g., despite ChEBI and MeSH both providing identifiers for small molecules, neither provides mappings to the other) and mappings that are provided are often incomplete (e.g. there are many gaps in mapping disease terms between MONDO and MeSH). Several automated methodologies have been proposed to predict missing mappings, including algorithms exploiting lexical similarity (Ghazvinian *et al*., 2009), ones based on logical/structural alignment of resources (Jiménez-Ruiz and Cuenca Grau, 2011), and others based on machine learning (Berrendorf *et al*., 2020). However, beyond routine benchmarking, mappings produced by such automated mapping algorithms are not systematically reviewed for correctness, and then contributed to primary sources to which the mappings apply, ultimately resulting in limited impact on the state of existing resources. Further, existing automated mapping approaches often do not provide interfaces for curating (e.g., reviewing, confirming, and rejecting) predicted mappings, storing important metadata (e.g., mapping confidence), and maintaining curation artifacts in accessible ways (e.g., via public version control). For example, the Ontology Alignment Evaluation Initiative (https://oaei.ontologymatching.org) has used the same task of aligning the Foundational Model of Anatomy Ontology (Rosse and Mejino, 2003) and SNOMED-CT (Donnelly *et al*., 2006) for more than fifteen years (see Bodenreider and Zhang (2006) and Wang and Hu (2022)). However, the manuscripts published on the task only focus on method development and neither predict novel mappings, curate them, nor contribute them back to the upstream identifier resources. Overall, existing work does not provide a workflow for finding, predicting, and curating missing mappings.

To address this, we introduce Biomappings (v0.2.0), a framework for semi-automatically creating and maintaining mappings in a public, version-controlled repository. Biomappings combines multiple contributions: (i) a “curation cycle” workflow for creating mappings, (ii) an extensible pipeline for automatically predicting missing mappings between resources, and automatically detecting inconsistencies, (iii) a web interface for reviewing and curating predicted mappings, and (iv) a public, version-controlled repository of predicted and curated mappings.

Biomappings currently makes available *∼*8.5 thousand reviewed mappings and *∼*41 thousand predicted ones. Mappings are associated with necessary metadata in an intuitive tab-delimited format; licensed permissively to encourage community contributions and restriction-free integration back into primary identifier resources. In addition to novel mappings, Biomappings provides an extensible lexical mapping prediction pipeline based on Gilda (Gyori *et al*., 2022), and a web-based interface for curation of predictions and adding manually constructed mappings. The mappings themselves, functions supporting the programmatic creation and usage of mappings, the web-based curation interface, as well as several workflow examples for generating new mappings are made available in the open-source *biomappings* Python package.

We demonstrate the utility of Biomappings in three case studies. First, we used Biomappings to predict and curate mappings for cell lines in the Cancer Cell Line Encyclopedia (CCLE) (Ghandi *et al*., 2019) to three other resources providing identifiers for cell lines. Biomappings added a total of 622 novel mappings that could not be inferred from existing mappings (a 71% increase). These mappings crucially improve the interoperability between databases describing the characteristics and measurements of cancer cell lines.

Second, we used Biomappings to predict and curate mappings between MeSH and ChEBI, both of which contain entries for chemicals of biological interest but lack any mappings to each others’ entries. We show that the 2, 663 mappings added by Biomappings enable mapping chemicals (listed using MeSH identifiers) in 100,265 ClinicalTrials.gov trials (70.5% of all trials) to ChEBI identifiers, thereby enabling the integration of clinical trials data with other ChEBI-aligned resources.

Third, we used Biomappings to predict and curate missing mappings between four widely used Open Biomedical Ontologies (OBO) (Jackson *et al*., 2021) ontologies and MeSH. We then contributed 1, 378 confirmed mappings back to the primary OBO resources through GitHub pull requests. These contributions have the potential for high impact given the ubiquity of these ontologies in various workflows, tools, and analyses.

Biomappings is available through a web portal (https://biopragmatics.github.io/biomappings) and as a Python software package (https://pypi.org/project/biomappings). All underlying data, code, and governance documentation are accessible through GitHub (https://github.com/biopragmatics/biomappings) under the MIT and CC0 licenses and are archived on Zenodo (Hoyt *et al*., 2022a).

## 2 Methods

### 2.1 The Biomappings curation cycle

We propose a novel, semi-automated approach to curation and maintenance of mappings supported by Biomappings (Fig. 1).

**Figure 1.**
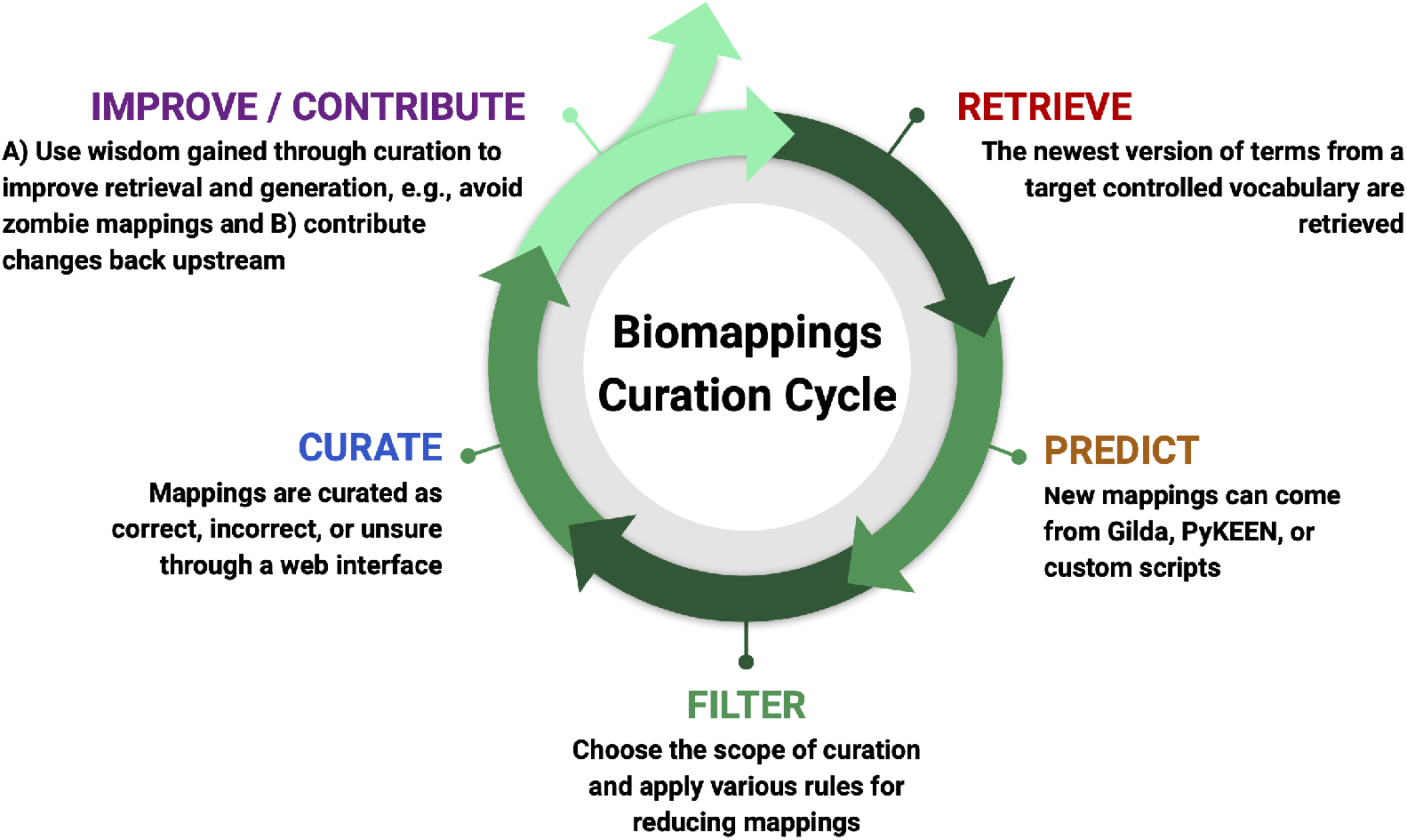
The Biomappings curation cycle supports sustainable, efficient curation of high-quality mappings between identifier resources, consisting of five steps.

The first step of the cycle comprises *retrieval* and preprocessing of target identifier resources, including any existing mappings between the resources. We automate this process for ontologies by using the Bioregistry (Hoyt *et al*., 2022c) to locate the ontology (i.e., with a URL) and ROBOT (Jackson *et al*., 2019) to parse it. Similarly, we use custom automated preprocessing workflows in PyOBO (Hoyt *et al*., 2022b) for other identifier resource types (e.g., databases like HGNC).

The second step comprises *generating predictions* through lexical similarity or other approaches (see Section 2.2). This step is often mediated by task-specific scripts which focus on generating mappings between two target identifier resources. At this stage, predictions for mappings that have already been manually curated in Biomappings or ones made available by other sources can be excluded to ensure predictions are novel.

The third step comprises choosing a scope for manual review, i.e., *filtering* predictions to a desired subset and applying additional filters. For instance, one may filter to predicted mappings between two specific identifier resources to focus curation on.

The fourth step is to review and *curate* predicted mappings as correct, incorrect, or unsure. In Biomappings, this step is mediated by a web interface described in Section 2.3. The fifth step involves *improving* the first four steps of the cycle based on insights gained from curation, as well as *contributing* curated mappings to external resources. For example, manual inspection of predictions may reveal systematic errors in prediction that can be used to improve preprocessing or update the filtering step in order to reduce curation burden.

Concurrently, the novel mappings can be contributed to increase their visibility and impact (we demonstrate this in Section 4.3).

This curation cycle workflow may support one of several goals such as 1) the exhaustive curation of mappings between identifiers in two identifier resources following previous rounds of curation, 2) a task-oriented curation goal to maximize return on fixed curation effort, often prioritized or driven by an external task-specific need, 3) analysis or quality assurance-driven curation such as the example described in Section 2.4.2, or 4) open-ended/interest-driven curation.

### 2.2 Generating Predictions

We employ a lexical matching workflow in the case studies presented in Section 4. This relies on labels and synonyms associated with entities for predicting mappings. The workflow is implemented using Gilda (Gyori *et al*., 2022), based on the text tagging algorithm from Allen *et al*. (2015). First, Gilda generates an index of the labels and synonyms for entities in the desired identifier resources by applying custom string transformations and normalizations (e.g., *Amyloid-β* becomes *amyloid-beta*). Second, Gilda applies the normalizations to each query string, identifies matches in the index, then calculates a score based on which normalizations were applied that resulted in the match (e.g., exact string matches have higher scores than strings with different usages of symbols or punctuation). For the lexical matching workflow, the labels and synonyms from each entity in a given identifier resource are used as the queries to an index built from the target identifier resource(s).

Lexical matching has the advantage of being computationally inexpensive, highly explainable, and ultimately easy to curate. However, we note that the Biomappings workflow is able to support other methodologies such as knowledge graph-based matching workflows (see Supplement), structural matching (e.g., that exploits ontology hierarchy, see Supplement), or other custom workflows. Following generation, predicted mappings are filtered to remove predictions that appear in primary identifier resources or have already been curated in Biomappings in order to reduce duplicate curation. Importantly, Biomappings also provides the framework for capturing provenance about how mappings are generated for all methodologies.

### 2.3 Web interface for curating mappings

Biomappings provides a locally-deployable web-based interface for browsing, reviewing, and curating predicted mappings (Fig 3). It displays a paginated view of the subject, predicate object, and confidence of each predicted mapping that can be searched by entity compact uniform resource identifier (CURIE), by entity name, and by resource to support restricted curation scopes. On the right-hand side, it displays buttons for curating each prediction as correct, incorrect, or unsure. After each curation, the interface updates the relevant resource files and includes provenance about the curation such as the curator’s ORCID identifier. It then communicates with `git` to create the appropriate commits that can be sent to GitHub. Finally, the bottom of the interface also includes a bar for inputting novel curations not included in the predictions.

**Figure 2.**
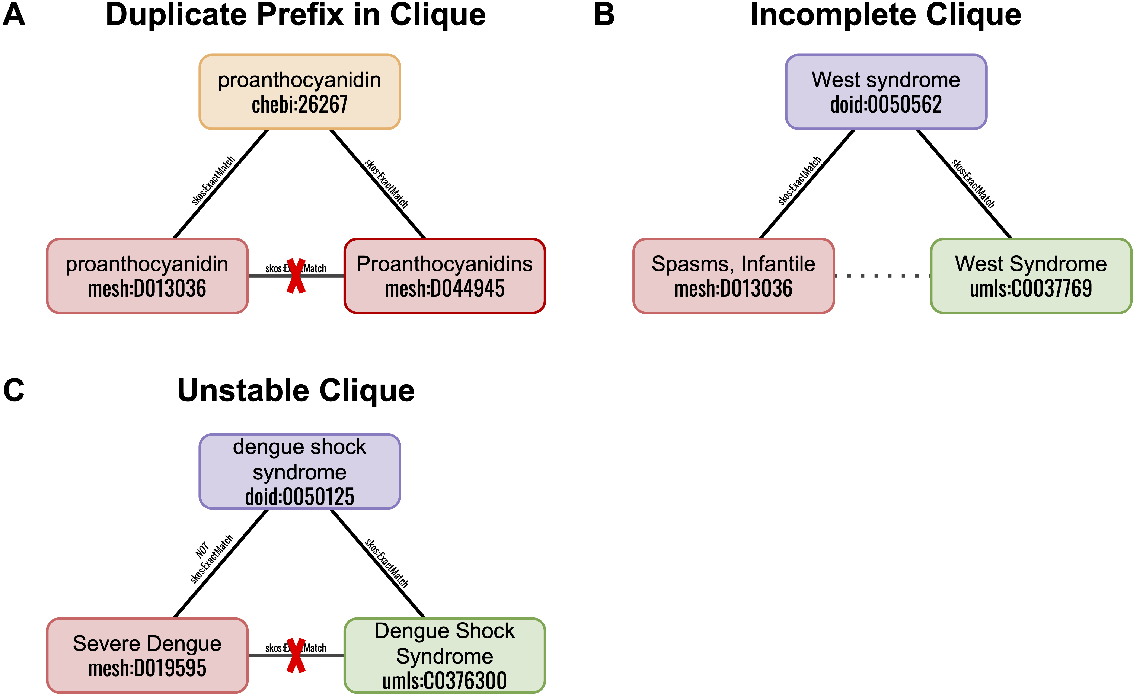
Three motifs identified by graph-theoretic methods for quality assurance. **A)** A prefix appearing twice in triangle of positive mappings signifies that one node and all of its edges is incorrectly mapped. **B)** A missing edge in a path of positive mappings with three nodes suggests the existing of a high confidence mapping. **C)** A triangle with a negative mapping and two positive mappings implies one of the positive mappings is incorrect.

**Figure 3.**
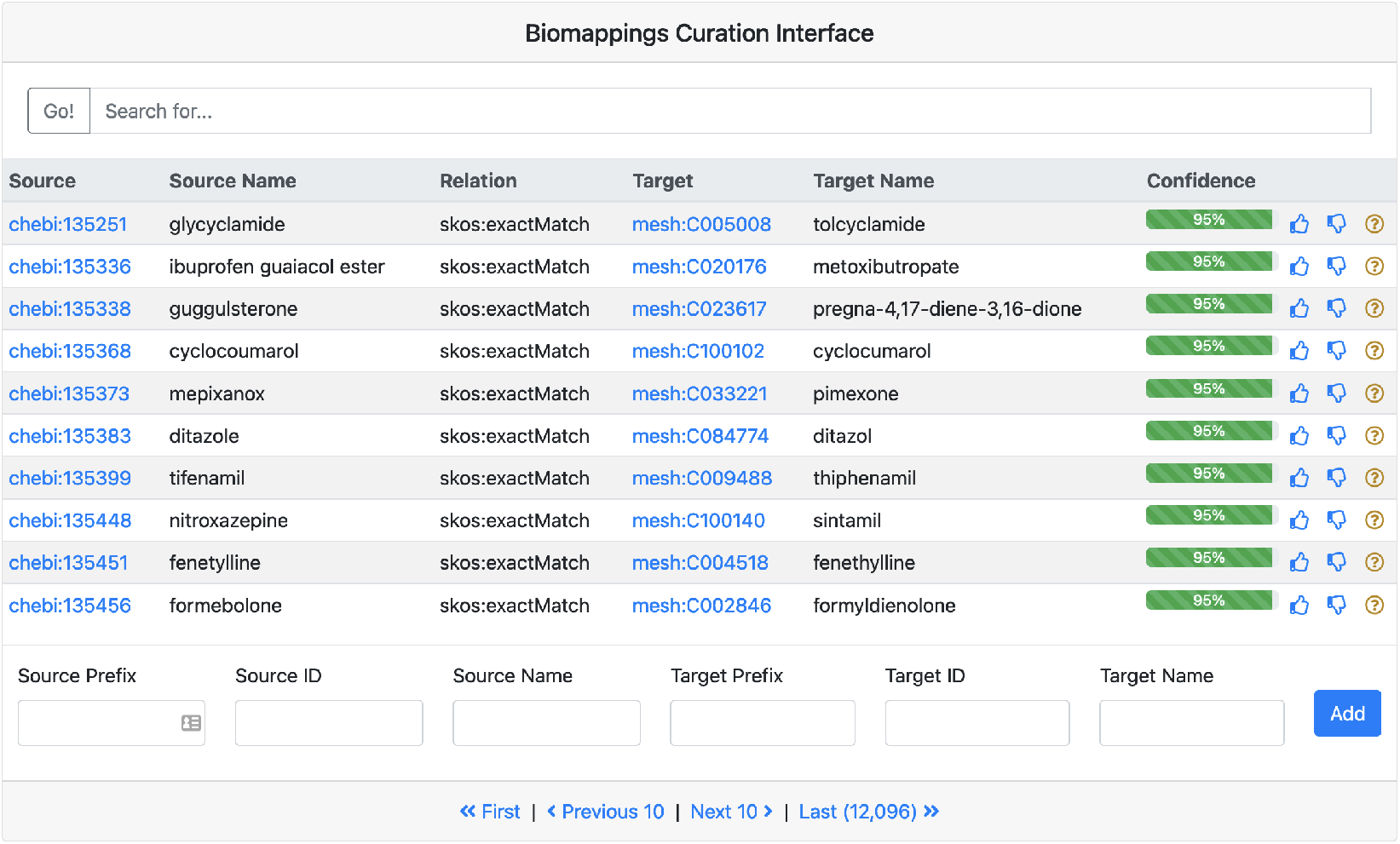
A screenshot of the Biomappings curation interface filtered to ChEBI and MeSH mappings showcases mappings of varying degrees of difficulty to curate. For example, some are exact matches, some are close matches, and some are matched due to synonyms.

### 2.4 Quality Assurance

Biomappings uses a combination of social and technical workflows to maintain high data quality and integrity.

#### 2.4.1 Version control and continuous testing

Biomappings uses `git` for version control to track all changes and mediate releases via Zenodo. Second, it uses GitHub as technical platform to host and distribute the project’s code and data openly and as a social platform to enable discussion and external contribution through pull requests. Third, it uses GitHub Actions as a continuous integration and continuous delivery system to apply data quality checks (e.g., all mappings’ prefixes and local unique identifiers are compliant with the Bioregistry (Hoyt *et al*., 2022c)). Further, several summaries (e.g., charts, tables, website), analyses (see Section 2.4.2), and artifacts (see Section 3.1) are automatically re-generated when a pull request is merged and archived on Zenodo (Hoyt *et al*., 2022a).

#### 2.4.2 Automated consistency checking of mappings

Biomappings implements three graph-theoretic approaches for identifying incorrect, inconsistent, or missing mappings. This serves as an automated quality check to maintain the global consistency of the resource.

First, a labeled, undirected graph is constructed from the union of positive and negative mappings. Then, certain pre-defined motifs are found as shown in Fig. 2 to detect inconsistencies.

The *duplicate prefix in clique* motif contains a set of nodes connected by equivalence relationships (e.g., `skos:exactMatch`) where two or more nodes originate from the same identifier resource (Fig. 2A). In the example, two entities from MeSH (i.e., *proanthocyanidin* (`mesh:C013221`) and *Proanthocyanidins* (`mesh:D044945`)) appear in the same clique. Upon further inspection, *Proanthocyanidins* was found to be represent a class of chemicals with similar structures and *proanthocyanidin* was found to represent a specific, prototypical instance of the class. Because the other entities in the clique referred to the specific chemical, *Proanthocyanidins* (i.e., the class) should be removed. Such motifs can also help identify more generic properties of the identifier resources that lead to curation errors, such as the pluralization schemes of its chemical classes.

The *incomplete clique* motif contains a set of nodes that are connected by equivalence relationships, but some nodes are not connected (Fig. 2B). This motif provides support for curating additional equivalence mappings via the transitivity of equivalence. In the example, a mapping between *Spasms, Infantile* (`mesh:D013036`) and *West Syndrome* (`umls:C0037769`) could be curated.

The *unstable clique* motif contains a set of nodes that are connected by equivalence relationships, but two of the nodes are also curated with a negative mapping (Fig. 2). This motif suggests that one (or more) of the equivalence or negative mappings are incorrect and should be removed. In the example, the equivalence between *Severe Dengue* (`mesh:D019595`) and *Dengue Shock Syndrome* (`umls:C0376300`) was incorrect.

These analyses are run on all changes to the Biomappings database using GitHub Actions as a continuous integration service and posts the results to https://biopragmatics.github.io/biomappings.

## 3 Results

Biomappings (v0.2.0) contains 8,560 positive mappings, 1,122 negative mappings, 47 unsure mappings, and 41,178 predicted mappings of 6 types across 27 identifier resources curated by 6 individuals (Table 1, see also detailed summary in Supplementary Fig. 2). These mappings were generated through a combination of prioritized curation (e.g., motivated by creating mappings to support a specific downstream task), open-ended curation (e.g., motivated by interest in a specific prefix or search term, or motivated by curating highest confidence mappings), and exhaustive curation (e.g., motivated by completing alignments between two or more identifier resources). Examples of each can be found in Section 4.

**Table 1.**
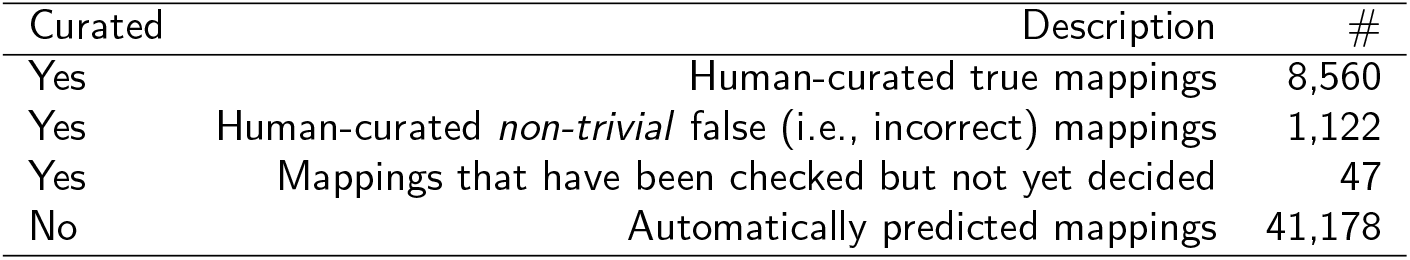
Biomappings resource files. Each row corresponds to a distinct tab separated value file in version control on GitHub.

| Curated | Description | # |
| --- | --- | --- |
| Yes | Human-curated true mappings | 8,560 |
| Yes | Human-curated *non-trivial* false (i.e., incorrect) mappings | 1,122 |
| Yes | Mappings that have been checked but not yet decided | 47 |
| No | Automatically predicted mappings | 41,178 |

While Biomappings is generally able to use any predicate encoded as a CURIE, mappings typically use the Simple Knowledge Organization System (SKOS) vocabulary to denote matches in the sense of information retrieval. The three most common predicates that are useful for curating mappings are `skos:exactMatch` for terms that can be used interchangeably, `skos:broadMatch` for when the object term is a super-class of the subject, and `skos:narrowMatch` for when the object term is a subclass of the subject. Three additional relations appear in Biomappings (v0.2.0) for species differentia (e.g., for mapping species-generic and species-specific pathways in KEGG (Kanehisa *et al*., 2017)), for homologs (e.g., for mapping related species-specific pathways in WikiPathways (Martens *et al*., 2021)), and for connecting different ionization states (e.g., conjugate acid, conjugate base) of small molecules (e.g., appearing in ChEBI).

### 3.1 Availability

Biomappings stores mappings in four tab-separated values (TSV) files corresponding to the correct, (non-trivially) incorrect, unsure, and predicted mappings (Table 1) that can be manually edited directly, modified via the back-end to web-based manual curation, or extended with new predictions programmatically. Each stores rich metadata about the source, target, mapping type, as well as its associated provenance. Further, the predicted mappings include confidence assessments and additional provenance about how they were generated. These are collated into a single SSSOM document that fully represents all provenance, can be used to generate additional exports (e.g., JSON, RDF), and can be readily consumed by curation workflows such as those in the Ontology Development Kit (ODK) (Matentzoglu *et al*., 2022b) for developing ontologies. These artifacts are re-generated by the continuous integration workflow triggered on all data changes in the repository. Further, these mappings can be accessed via the `biomappings` Python package, installable through the Python Package Index (PyPI). Finally, the mappings are available as an interactive network on the Network Data Exchange (NDEx) (Pratt *et al*., 2015) under https://bioregistry.io/ndex:402d1fd6-49d6-11eb-9e72-0ac135e8bacf.

### 3.2 Governance and Sustainability

Biomappings is built using open source code and open data distributed under a permissive license (MIT and CC0) in a public, version-controlled repository that is archived on Zenodo in order to encourage community reuse and incorporation into upstream resources that it describes. It leverages public infrastructure and automation to support its maintenance and extension. It has well-defined contribution guidelines (https://github.com/biopragmatics/biomappings/blob/master/docs/CONTRIBUTING.md) and a governance model (https://github.com/biopragmatics/biomappings/blob/master/docs/GOVERNANCE.md) that enables contributions directly from the broader community to support the project’s longevity. Finally, Biomappings has a transparent attribution model that associates all mappings with the Open Researcher and Contributor Identifier (ORCID) of the curator. These are summarized on the auto-generated summary website and also on APICURON (Hatos *et al*., 2021).

## 4 Case Studies

We present three case studies demonstrating the utility of Biomappings. In Section 4.1, we describe generating and curating exhaustive mappings between several identifier resources for cancer cell lines. In Section 4.2, we describe taking a prioritized approach towards generating and curating mappings between ChEBI and MeSH in order to support data integration with clinical trials data. Finally, in Section 4.3, we describe predicting and curating missing MeSH mappings for several OBO ontologies, then contribute the results back to the primary identifier resources.

### 4.1 Mapping missing cancer cell line identifiers

Several identifier resources have been constructed to describe cells and cell lines in order to support the generation of databases describing their characteristics and associated experimental measurements. For example, the Cancer Cell Line Encyclopedia (CCLE) contains a detailed genetic and pharmacological characterization of hundreds of cancer cell lines constituting models of various human cancers. Such detailed, large-scale databases have proven useful in the prediction of anti-cancer drug sensitivity (Barretina *et al*., 2012) and pre-clinical testing Wilding and Bodmer (2014).

We were interested to integrate experimental data from CCLE in a dialogue system, but because CCLE provides an identifier resource that does not contain lexicalizations (i.e., names and synonyms), it was necessary to map CCLE to external identifier resources in order to use their respective lexicalizations. Cellosaurus (Bairoch, 2018) and the Experimental Factor Ontology (EFO) (Malone *et al*., 2010) were chosen because of their high quality curation, detailed lexicalizations, and pre-existing mappings to CCLE and related identifier resources.

EFO (v3.47.0) contains no direct mappings to CCLE, but Cellosaurus (v43.0) contains 1,448 (black lines in Fig. 4). Cellosaurus also contains 1,302 mappings to EFO which theoretically enabling the inference of 723 two-hop mappings from CCLE to EFO (blue dotted line in Fig 4). Further, we were able to recover a partially overlapping three-hop mapping of 725 from CCLE to EFO when using the Cancer Dependency Map (DepMap) as an intermediate (red dotted line in Fig 4). However, such inference is inconvenient, relies on several implicit assumptions, and is still incomplete.

**Figure 4.**
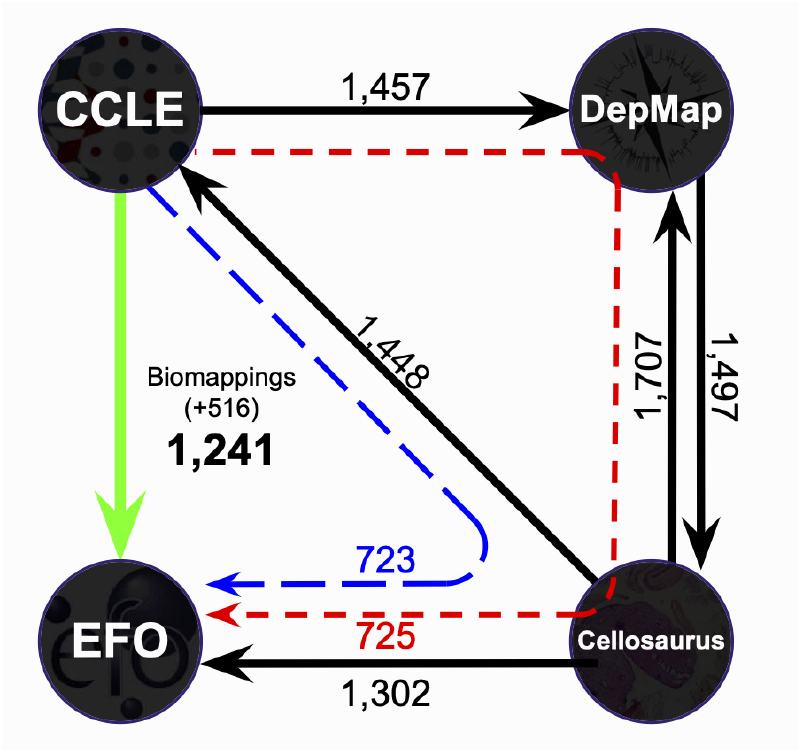
Mappings between identifiers in four cancer cell line resources. Solid black arrows represent primary mappings provided by identifier resources (arrows point from an identifier resource that provides a mappings to its target, labels show the number of mappings provided). Dashed arrows represent inferrable mappings through two hops (blue, 723 mappings inferred) or three hops (red, 725 mappings inferred). The green arrow represents the added benefit of Biomappings curation of mappings between CCLE and EFO, 1,241 total with 516 not obtainable from existing primary or inferred mappings.

In order to complete the alignments between these three identifier resources, we generated novel lexical mappings (i.e., that were not inferrable) and curated 106 (+7%) from CCLE to Cellosaurus and 516 (+71%) from CCLE to EFO. We also curated 30 nontrivial negative mappings from CCLE to Cellosaurus and 9 non-trivial negative mappings from CCLE to EFO to help avoid future mismatches. For example, we asserted that *CL14_LARGE_INTESTINE* (`ccle:CL14_LARGE_INTESTINE`), a colorectal adenocarcinoma, is not the same as *XPCS2BASV Cl-14* (`cellosaurus:ZP40`), a transformed fibroblast.

While this case study focused on aligning cancer cell line terms, it represents an important first step in more generally improving the interoperability between cell and cell line resources that support comparative analysis, generation and characterization of disease models, and many other efforts in drug discovery.

### 4.2 Chemical identifier mappings to improve clinical trials data integration

ClinicalTrials.gov is a database of publicly funded clinical studies provided by the United States National Library of Medicine. It provides bulk programmatic access that includes for each trial which include the MeSH terms for around 3,600 unique interventions (e.g., small molecules) and around 4,200 unique conditions. However, MeSH does not provide primary mappings to highly accessible identifier resources (e.g., ChEBI) that are necessary to support data integration with other popular data sets using different identifier resources^1^.

In this case study, we focused on generating and curating lexical mappings from MeSH to ChEBI. Many interventions appear across multiple clinical trials, so rather than exhaustively curating MeSH term mappings for all unmapped interventions, we prioritized curation by frequency of appearance across all 430K clinical trials. Because the distributions of frequencies of interventions has a long tail, we selected a subset of MeSH terms that represented 80% of the respective unmapped trial-intervention instances.

This resulted in the curation of mappings from 282 interventions’ MeSH terms to ChEBI that covered 80% (120K) of the 150K trial-intervention pairs with unmapped interventions. Note that some previous curations from MeSH to ChEBI already existed in Biomappings from a combination of undirected curation and various task-specific curations, causing this number to be lower than if we had conducted this curation at the beginning of the Biomappings project. Further, iterations of identifying the most valuable 80% of remaining unmapped interventions becomes successively less impactful due to the remaining interventions appearing in fewer trials.

Ultimately, Biomappings (v0.2.0) contains 2,663 mappings from MeSH to ChEBI which enables mapping 100,265 (70.5%) clinical trials to 987 unique chemicals. These mappings enable the previous difficult integration of clinical trials data from ClinicalTrials.gov with other resources using (or mappable to) ChEBI. For example, a knowledge graph containing ClinicalTrials.gov, chemical-protein bioactivities in ChEMBL (Gaulton *et al*., 2017), and protein-pathway memberships in WikiPathways can be used to answer questions like *“What are the protein targets modulated in a given clinical trial?”* and *“What are the pathways modulated in a given clinical trial?”* with simple graph queries relying on.

### 4.3 Extending community ontologies with missing mappings

Several high quality OBO ontologies cover similar topics/domains as MeSH and therefore maintain mappings to relevant MeSH terms. When reviewing mappings to MeSH in multiple widely used community ontologies, we found that mappings were incomplete. We therefore predicted (using an automated lexical approach) and predicted mappings from four OBO ontologies to MeSH outlined in Table 2.

**Table 2.** A summary of upstream contributions of 1,378 Medical Subject Headings (MeSH) mappings back to primary ontology resources

| Resource | # | Link |
| --- | --- | --- |
| Human Disease Ontology (DO) | 974 | PR #1073 |
| Monarch Disease Ontology (MONDO) | 190 | PR #4930 |
| UBERON (Haendel *et al.*, 2014) | 130 | PR #2432 |
| Cell Ontology (CL) (Diehl *et al.*, 2016) | 84 | PR #1561 |

We then implemented automated scripts for inserting the new mappings into the respective version-controlled source files for each ontology in the OWL/XML, functional OWL, or OBO text formats, depending on the ontology. Finally, we made pull requests on the associated GitHub repository for each ontology to integrate the proposed mappings made using Biomappings through which ontology maintainers were able to add these contributions. Contributing curated mappings upstream is important and highly impactful because it is propagated directly to users, generic services that consume ontologies such as Ubergraph (Balhoff and Curtis, 2021), other services that consume mappings such as the Ontology Mapping Service (EMBL-EBI, 2022), and pipelines that build knowledge graphs (e.g., Phenotype Knowledge Translator (PheKnowLator) (Callahan, 2019)). Together, these form the basis for a large number of computational workflows used across by researchers and engineers in academic, industrial, and research institutions.

## 5 Discussion

We presented Biomappings, a repository for community curated mappings between biomedical entities. Biomappings stores predictions and manual curations along with granular metadata and high-level semantics for each. It relies on an open data, open code, open infrastructure philosophy combined with a novel governance strategy to foster community contributions and engagement. It also explicitly encourages reuse and redistribution via its highly permissive CC0 license. Biomappings uses public infrastructure for quality assurance and distribution to promote transparency and increase trust.

### 5.1 Limitations

Biomappings enables the curation of missing mappings that are not available from primary identifier resources. In some cases, however, identifier resources have idiosyncratic curation guidelines for what constitutes a mapping, e.g., in how strictly two terms need to match to justify a mapping. This means that conflicts may arise if the curators of a primary identifier resource differ in their interpretation with what is curated via Biomappings. These can be resolved through community engagement and discussion, and - as demonstrated in this paper - direct contribution to primary resources in a public space.

Further, even after completing an exhaustive curation campaign to create mappings between two identifier resources, it is still possible that new terms will be added late, even more complex operations such as the splitting or merging of terms, changes to the scope of a term’s definition. The automation of predictions in Biomappings can facilitate keeping up with updates to identifier resources, however, automation does not yet extend to interpreting more complex changes to resource over time, requiring manual review.

### 5.2 Extensibility

The initial focus of the Biomappings project is on entities in the biomedical domain. However, we believe that the tooling and philosophy can be readily adapted to domains outside of biomedicine, in fields where multiple overlapping identifier resources exist and need to be mapped for data integration. One such example is agriculture and agronomy: as a proof of concept, Biomappings includes 142 curations contributed to align concepts related to soil in the Agronomy Ontology (Arnaud *et al*., 2020) and Agronomy Vocabulary (Mietzsch *et al*., 2021). This demonstrates the possibility of the extension of Biomappings’ scope and the project’s ability to motivate external contribution.

In our case studies, we used a lexical approach to predict missing mappings between resources. However, the Biomappings curation cycle does not depend on the specific approach to two resources are aligned to identify potential mappings. For instance, knowledge graph-based or machine learning-based methods such as those demonstrated by Berrendorf *et al*. (2020) can be readily used with Biomappings and our future work will involve making use of such methods to find mappings missed via lexical alignment.

### 5.3 Future Work on Biomappings

Following the initial development and curation of Biomappings, two ongoing challenges remain. First, there are multiple ways in which the (semi-)automated approach to the generation and curation of predictions could be improved in the future. These include improving the data model to propagate information about the version of each identifier resource used for the generation of predictions, and the ability to re-run generation workflows in an automated way, then to notify relevant curators that new content is available. In order to reduce curation burden, maximize return on curator time, and ultimately to scale curation of mappings, we will need to develop more accurate approaches for prediction, filtering, and prioritization.

The second challenge is to build, train, and engage a community of curators. One sub-challenge of this is to create curation interfaces that can be more easily deployed (or hosted) and seamlessly interact with git and GitHub (e.g. similar to the OntoDev suite, see https://github.com/ontodev) to support potential curators who might not be familiar with common software workflows or comfortable with version control. A second sub-challenge is to improve the ability to contribute content curated in Biomappings back to primary identifier resources. While we demonstrated this for four OBO ontologies, there are both technical challenges (e.g., the data is not curated in public version control) and social challenges (e.g., the maintainers are not receptive to contribution) for contributing content to primary resources. Distributing the burden of curation, for example, using Biomappings, has substantial potential to improve ontologies which are used either directly or indirectly by most modern scientists.

### 5.4 Future Vision

We envision the broader community of curators and developers who create and consume ontology mappings working in several directions to better support the tasks (e.g., ontology merging, entity alignment) that rely on mappings. First, we hope to see more resources adopting formats and minimum metadata standards like SSSOM to make their mappings more reusable (e.g. by assigning more precise predicates and including provenance information). Second, we hope to see these resources converging on external standards for the syntax and semantics used to communicate the entities and predicates appearing in mappings, such as the Bioregistry (Hoyt *et al*., 2022c) in order to improve interoperability. Third, we hope to see large-scale efforts to aggregate, store, and redistribute mappings with more general scope than existing mapping services. Finally, we hope to see the development, implementation, and deployment of standardized, efficient algorithms for inference and retrieval of mappings as well as an associated provenance model. We believe Biomappings will have an important role in supporting curation efforts and applications of mappings as the community works towards these goals.

## Acknowledgements

We acknowledge Daniel Domingo-Fernández and Sarah Mubeen for contributing mappings from PathMe and Krishna Udaiwal for contributing agronomy mappings. We would also like to thank Harry Caufield for the helpful discussions.

## Footnotes

1 Some mappings are available from MeSH to CAS and UNII, but these are not annotated well and these resources are relatively inconvenient to use

