## Supplementary information for "Prediction and Curation of Missing Biomedical Identifier Mappings with Biomappings"

### Biomappings Supplementary Information

#### Table of contents

##### Supplementary sections

Supplementary Section 1. Structure and summary of mappings.
Supplementary Section 2. Additional matching methodologies.

##### Supplementary figures

Supplementary Figure 1. Structure of manually curated mappings table.
Supplementary Figure 2. Summary of positive manually curated mappings.

#### 1 Structure and summary of mappings

Biomappings contains four tab-separated values (TSV) files for predicted mappings and positive, negative or unsure curated mappings. Each row in these files represents a single mapping. Each mapping represents a subject (i.e., source) and a target (i.e., object) for the mapping with a prefix, local unique identifier, and standard name for each in separate columns. Storing the prefix and the identifier in separate columns allows for convenient filtering by prefix in downstream usage, while storing the standard name - though redundant - facilitates human interpretability of mappings. In addition, each mapping represents a specific relation (i.e., predicate) as a compact URI (CURIE). Currently used predicates include `skos:exactMatch` for exact equivalence - this is the most common predicate in the current curated set - and `skos:narrowMatch` or `skos:broadMatch` for non-exact matches. Further, we use two other predicates for special relationships: `speciesSpecific` for relations where a subject is not an exact match but rather a species-specific variant of a non-species-specific object, and `RO:HOM0000017` ("in orthology relationship with") to represent cross-species orthologs of the same gene/protein. Biomappings is not limited to these predicates though, and future mappings can use other predicates as needed.

Predicted and curated mappings differ in what provenance columns are included. For curated mappings, each row includes a "type" to denote whether a mapping was predicted and then reviewed or manually curated *de novo*, as well as the curator's ORCID identifier. For predicted mappings a "type" column represents the method by which the mapping was predicted, e.g., "lexical", and a "source" column provides the URL to the script that created the mapping. The predictions file also includes a confidence score between 0 and 1, where a higher score represents a prediction more likely to be correct. When producing predictions, scores are ideally set to approximate the empirical precision associated with the predictions.

The Biomappings files can be accessed as follows:

1. Predicted mappings `https://github.com/biopragmatics/biomappings/blob/master/src/biomappings/resources/predictions.tsv`
2. Curated correct mappings `https://github.com/biopragmatics/biomappings/blob/master/src/biomappings/resources/mappings.tsv`
3. Curated incorrect mappings `https://github.com/biopragmatics/biomappings/blob/master/src/biomappings/resources/incorrect.tsv`
4. Curated unsure mappings `https://github.com/biopragmatics/biomappings/blob/master/src/biomappings/resources/unsure.tsv`

Figure 1 shows several exemplar rows from the manually curated positive mappings file.

Figure 2 provides a more detailed breakdown of which identifier resources have been covered by Biomappings curation. Medical Subject Headings (MeSH) Rogers (1963) appears on top, due to the fact that it doesn't provide mappings to several key external resources, therefore a large portion of the curation effort in Biomappings has focused on mapping MeSH to various chemical, protein, anatomical, and disease identifier resources.

| | source prefix | source identifier | source name | relation | target prefix | target identifier | target name | type | source |
|---|---|---|---|---|---|---|---|---|---|
| 1 | ccle | 143B_BONE | 143B | skos:exactMatch | cellosaurus | CVCL_2270 | 143B | manually_reviewed | orcid:0000-0003-1307-2508 |
| 2 | ccle | 143B_BONE | 143B | skos:exactMatch | efo | 0006355 | 143B | manually_reviewed | orcid:0000-0003-1307-2508 |
| 3 | ccle | 59M_OVARY | 59M | skos:exactMatch | cellosaurus | CVCL_2291 | 59M | manually_reviewed | orcid:0000-0003-1307-2508 |

Figure 1: A screenshot of the manually curated mappings table in the Biomappings version-controlled repository. This contains information about the source entity, the target entity, the predicate, the type of mapping, and the curator.

### 2 Additional Matching Methodologies

**Rule-based Lexical Matching Methods** Rule-based lexical matching methods rely on deterministic, data source-specific rules for generating high confidence predicted mappings. For example, because the MeSH supplement contains terms for human proteins whose names are generated based on the labels in UniProt Bateman *et al.* (2021), mappings can be generated with deterministic text processing and matching (e.g., *AKT1 protein, human* (`mesh:C494918`) maps to *AKT1* (`uniprot:P31749`)). Similarly, WikiPathways Martens *et al.* (2021) labels for homologous pathways can be used to generate exact string matches after stripping the organism ellipses (e.g., *Apoptosis (Homo sapiens)* (`wikipathways:WP254`) maps to *Apoptosis (Mus musculus)* (`wikipathways:WP1254`)).

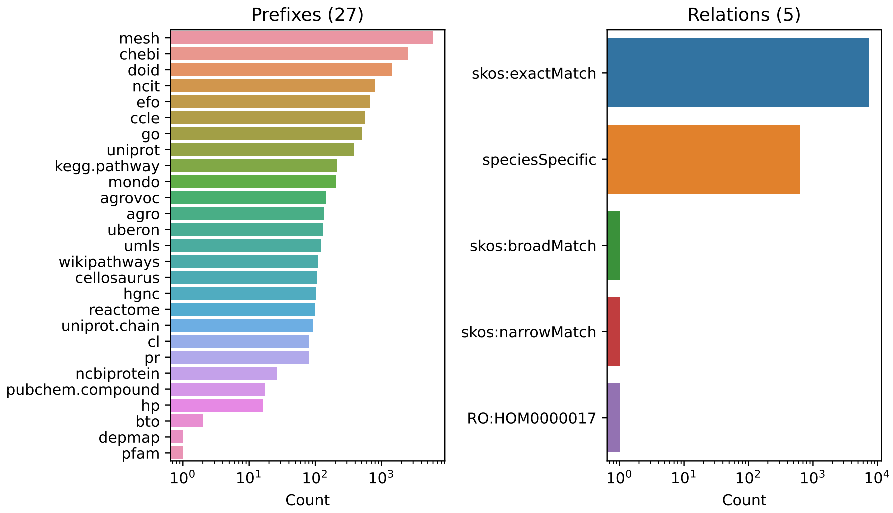


Figure 2: A summary of curations in Biomappings, broken down on the left by prefix and on the right by relation type.

**Structural Matching Methods** Structural matching draw from a combination of local properties of entities in identifier resources (e.g., their properties and relationships) as well as global properties (e.g., taxonomical structure, graph structure) to infer or predict mappings (Chauhan *et al.*, 2018). For example, graph homomorphism-based methods search for sub-graphs with similar structure that can be used to infer mappings. (Chauhan *et al.*, 2018). Constraint-based methods have a similar approach but alternate mathematical formulations (Mao *et al.*, 2010; Algergawy *et al.*, 2008). Taxonomy-based matching algorithms focus on hierarchical information for matching (Nandi and Bernstein, 2009) while semantic similarity methods include additional features, e.g., from textual labels or descriptions (Essayeh and Abed, 2015; Wang *et al.*, 2010; Xiang *et al.*, 2015) that are featurized in various ways. Several of these methods rely on machine learning approaches for inference while others are more classical procedural algorithms.

**Knowledge Graph-based Matching Methods** Knowledge graph-based entity alignment methods incorporate prior knowledge and data about concepts to predict mappings, often with knowledge graph embedding models Berrendorf *et al.* (2020). These methods construct disjoint knowledge graphs with concepts and relations from each respective controlled vocabulary (typically ontologies), connect the graphs with triples corresponding to both primary and third-party mappings, then train a link prediction model to predict novel mappings. While these models promise to generate more interesting predicted mappings, they have the drawback that they are slower to train and lack explainability.
